# FabriEVEs: A dedicated platform for endogenous viral elements in fishes, amphibians, birds, reptiles and invertebrates

**DOI:** 10.1101/529354

**Authors:** Jixing Zhong, Jiacheng Zhu, Zaoxu Xu, Geng Zou, JunWei Zhao, SanJie Jiang, Wei Zhang, Jun Xia, Lin Yang, Fang Li, Ya Gao, Fang Chen, Yiquan Wu, Dongsheng Chen

## Abstract

Endogenous viral elements (EVEs) are heritable viral deriving elements present in the genomes of other species. As DNA ‘fossils’ left by ancient viruses, EVEs were used to infer the characteristics of extinct viruses. EVEs in mammals have been well classified by several databases, however, EVEs in non-mammalian organisms are poorly documented. Here, we report FabriEVEs (http://tfbsbank.co.uk/FabriEVEs), the first dedicated and comprehensive online platform for the collection, classification and annotation of EVEs in fishes, amphibians, birds, reptiles and invertebrates. In total, nearly 1.5 million EVEs from 82 species deriving from class I (dsDNA), II (ssDNA), III (dsRNA), IV (positive ssRNA), V (negative ssRNA), VI (ssRNA-RT) and VII (dsDNA-RT) viruses were recorded in FabriEVEs, accompanying with comprehensive annotation including the species name, location, genomic features, virus family and associated literature. Flexible and powerful query options were provided to pinpoint desired EVEs. Furthermore, FabriEVEs provides free access to all EVEs data in case users need to download them for further analysis. Taken together, our database provided systematic classification and annotation of EVEs in non-mammal species, which paves the way for comparative analysis of EVEs and throws light upon the co-evolution of EVEs and their hosts.

## INTRODUCTION

Although endogenous viral elements (EVEs) mainly origin from retroviruses(1–4), increasing number of EVEs have been identified from non-retroviruses families including parvoviruses, filoviruses, bornaviruses and circoviruses(2). EVEs can be classified into partial EVEs or entire EVEs (proviruses) depending on whether they represent the whole genomes of extant viruses. EVEs provide a unique opportunity for researchers to investigate extinct viruses, which give rise to the emerging Paleovirology(5). Comparison EVEs with their extant counterparts give us a clue regarding the evolutional trajectories of viruses and the co-evolution or the arm race between viruses and hosts(2,6). EVEs have been found to play important roles in regulating the expression of host genes by introducing novel cis-regulatory elements including promoters and enhancers(1,7–9). Domesticated EVEs are found to be involved in several critical biological processes of hosts such as immunity(10) and the formation of placenta(11,12). The contribution of immunity by EVEs can be interpreted in two aspects. Firstly, EVEs were found to regulate several immune-related cellular genes and were essential for the innate immunity of host(13). Besides, Some endogenous viral genes can be domesticated and utilized by hosts to fight against exogenous viruses by blocking their cell entry and replication(10,14–18).The initial studies of EVEs mainly focused on mammals, probably attributed to the available integral and abundant whole genome sequences. Recently, increasingly more EVEs have been identified in non-mammals species, along with the progress of modENCODE(19), Genome10K program(20), Avian Phylogenomics Project(21) and B10K Project(22). In fishes, several cases of EVEs have been reported in spotted green pufferfish and elephant shark respectively(23,24). A systematic screen on 48 bird species revealed a variety of EVEs from both retroviruses and non-retroviruses, though the frequency of EVEs is much lower than that in mammals(4). In reptiles, EVEs deriving from hepadnaviruses, bornaviruses and circoviruses have been identified from several snake species and Chinese soft shell turtle(25,26). EVEs have also been found in amphibians such as western clawed frog (*Xenopus tropicalis*)(27). In invertebrates, 27 species were found to accommodate several virus origin fragments(6,26,28). Of particular interest, a remarkable diversity of EVEs was found to exist in a crustacean genome, pill bug (*Armadillidium vulgare*)(29). Up to now, there are two EVEs databases for mammals. HERVd, the first database for human endogenous retroviruses, was released in 2002 and updated in 2004(30,31). gEVE version 1.1 houses 774,172 EVEs identified from 20 mammalian genomes(32). Despite a large amount of EVEs have been identified and reported in non-mammal species, there is no specific database to accommodate these invaluable data sets. To fill the gap between increasing EVEs data in fishes, amphibians, birds, reptiles and invertebrates and our limited ability to interpret them, we developed a dedicated online platform for the storage, classification, annotation, querying and downloading of non-mammal animal EVEs (Figure 1).

**Figure 1.**
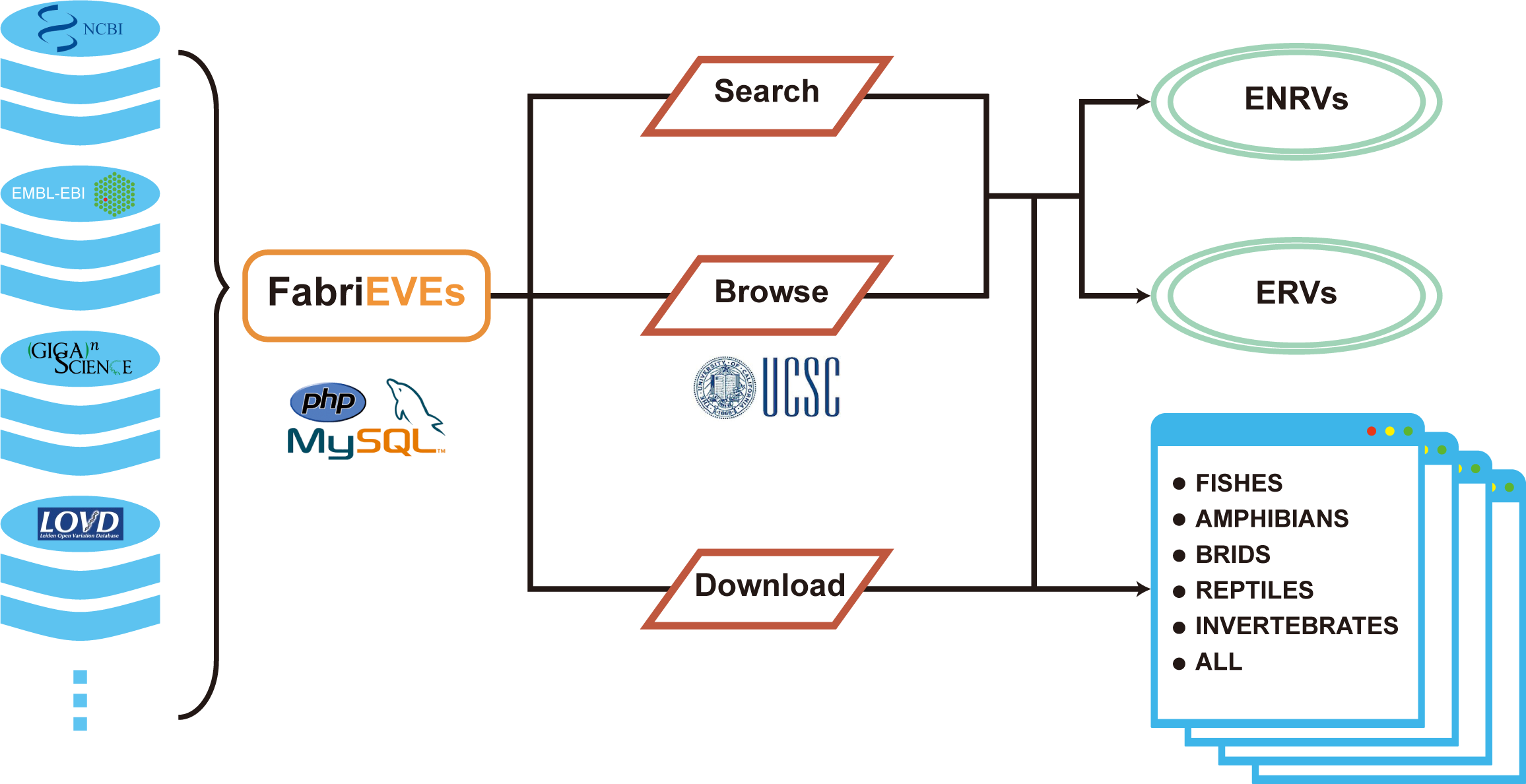

### The statistics of data in FabriEVEs

In total, 1,483,658 EVEs from 19, 1, 48, 4 and 28 species in fishes, amphibians, birds, reptiles and invertebrates were included in FabriEVEs (Table 1). Although the majority of the EVEs (>99.99%) derived from retroviruses as expected, we still observed a high diversity of EVE originating from non-retroviruses. For example, in birds, 249 EVEs from hepadnaviruses (double strand DNA viruses), bornaviruses (negative single strand RNA viruses), circoviruses (single strand DNA viruses) and parvoviruses (single strand DNA viruses) were presented in our database. It is of particular interest that some species accommodate much more EVEs than their close relatives. In birds, *Phalacrocorax carbo* (Cormorant) contains 68 copies of non-retroviruses EVEs, compared to 0-14 copies of non-retroviruses EVEs in remaining 47 bird species. Admittedly, this unusual high copy number of non-retroviruses EVEs in Cormorant might simply be due to differences in genome size or genome assembly among the 48 sequenced bird species. However, we ruled out this possibility based on following reasons: 1) All of the genome size range from 1.05 Gb in *Acanthisitta chloris* (Rifleman) to 1.26 Gb in *Aptenodytes forsteri* (Emperor penguin), which means that no bird genome is significantly larger than the others among the 48 sequenced genomes in the Avian phylogenomics project(33–36). 2) The sequencing depth of Great cormorant is only 24X compared to the depth of 160X in *Melopsittacus undulatus* (Budgerigar); 3) Among those 48 species, *Gallus gallus* (chicken) is the most well-studied bird species with its draft genome released as early as 2004(37). However, no non-retroviruses EVEs have been reported in chicken genome according to Cui’s systematic screen on the 48 bird species(4). In summary, the high copy number of non-retroviruses EVEs in *Phalacrocorax carbo* (Cormorant) was unlikely to be caused by genome size or genome assembly variance. Considering previous reports have shown that EVEs might play important roles in the evolution of animals(1–3), it is reasonable to speculate that those *Phalacrocorax carbo* (Cormorant)-specific non-retroviruses might contribute to its divergence from other species or facilitate its adaptation to its unique ecology niches.

**Table 1.** Number of EVEs in fishes, amphibians, birds, reptiles and invertebrates deriving from several viruses families(*hepadnaviridae, bornaviridae, circoviridae, parvoviridae, filoviridae and retroviridae*).

| Class | Hepadnaviridae | Bornaviridae | Circoviridae | Parvoviridae | Filoviridae | Retroviridae | Others | Total |
| --- | --- | --- | --- | --- | --- | --- | --- | --- |
| Fishes | 0 | 0 | 0 | 2 | 0 | 19 | 0 | 21 |
| Amphibians | 0 | 0 | 0 | 0 | 0 | 1 | 0 | 1 |
| Birds | 225 | 6 | 5 | 13 | 0 | 1,478,857 | 0 | 1,479,106 |
| Reptiles | 4 | 3 | 6 | 0 | 0 | 0 | 2 | 15 |
| Invertebrates | 0 | 1 | 20 | 12 | 28 | 4183 | 269 | 4513 |

### EVEs data Search

As there are 1.5 million EVEs in FabriEVEs, we provided advanced methods for users to retrieve the data they are looking for (Figure 2a). Specifically, four key attributes of each EVE elements were used to build the query. To begin with, the first filtering method was based on the class and scientific name of species. Class names (fishes, amphibians, birds, reptiles and invertebrates) were used to initiate the classification. After choosing the specific class, all the species in this class will be shown in a selection list, from which users can choose what species they are interested in. In addition, EVEs can be classified into *hepadnaviridae, bornaviridae, circoviridae, parvoviridae, densovirinae, reoviridae, rhabdoviridae, flaviviridae, bunyaviridae, orthomyxoviridae, retroviridae* or “unassigned” based on which family the corresponding extant counterpart viruses belong to. All those options were provided as check-boxes and the default status for them are ‘checked’. Users can deselect singular or several of them, and the corresponding types of EVEs will be included or excluded from the results. Similarly, EVEs can be divided into partial or entire depending on whether they are homologous to part or entire genome sequences of extant viruses. After choosing desired options for class, species, type and completeness of EVEs, all the selected values will be submitted to a PHP script, which will execute the query based on provided information. For example, if the class ‘bird’, species ‘*Acanthisitta chloris*’, EVE type ‘hepadnaviruses’, completeness ‘entire’ were chosen, then only 1 record fulfilling all those criteria will be returned. That is ‘Rifleman_hepadnaviruses_1’ located on the scaffold27280 of *Acanthisitta chloris* (Rifleman). If multiple records were found, then they will be listed in a table (Figure 2e). Each row begins with a unique EVE ID such as Rifleman_hepadnaviruses_1, which contains the species name, EVE type and a number to indicate its rank in the database. To provide more detailed information for every EVE, the EVE ID was hyperlinked to a detail page.

**Figure 2.**
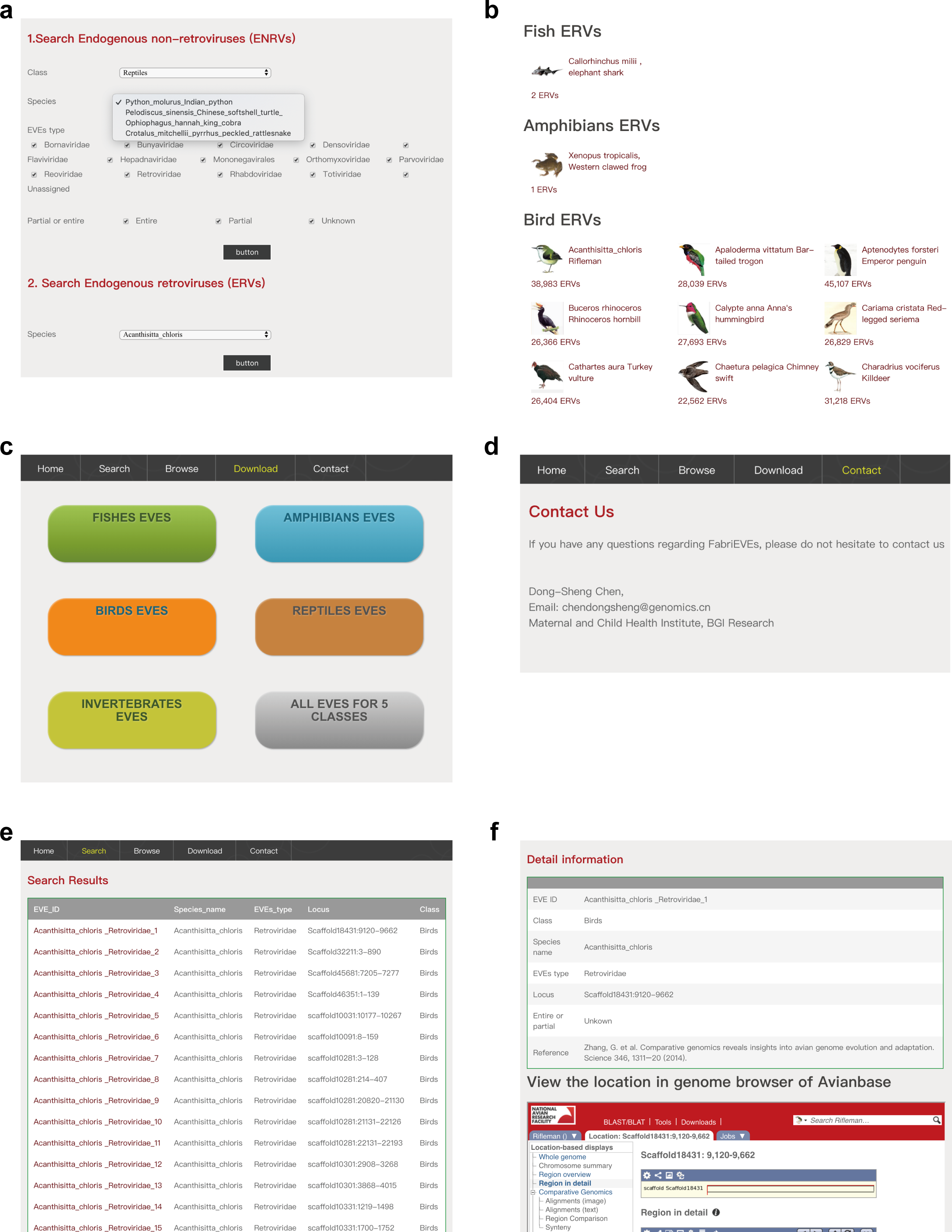

### EVEs detail demonstration

In the detail information page, the following information was provided (Figure 2f): 1) EVE ID indicating the unique identifier of the EVE in FabriEVEs database; 2) Class, common name and species name specifying the name of certain animal species; 3) EVEs type representing the type of EVEs; 4) Locus showing the location of EVE on animal chromosomes or contigs; 5) completeness indicating whether the EVE is complete or partial; 6) Reference showing from which literature the EVEs data is retrieved from;

### EVE data downloading

In addition to comprehensive annotation, we provided free access to all the EVEs in our database (Figure 2c). In the downloading page, EVEs data from fishes, amphibians, birds, reptiles and invertebrates were compressed as zip format files and hyperlinked to a button with the information of host class and the number of EVEs in that corresponding class. Therefore, users can download data sets from any class by clicking that button. These datasets are also available in CNGBdb.

### Analysis of functions of EVEs in Drosophila melanogaster

To explore the functional implications EVEs, we performed comprehensive analysis on the EVEs elements in Drosophila melanogaster. In total, 4183 Drosophila melanogaster EVEs elements were stored in FabriEVEs and but after filtering only 3594 of them were analyzed using ChIPseeker (references). The results suggest that the majority of EVEs were distributed on chromosome 2R, 3L and 3R (Figure 3a). To assess the genomic features of EVEs, we calculated the frequency of EVEs in different genomic regions including promoter (<=1kb), promoter (1-2kb), promoter (2-3kb), 5’ UTR, 3’ UTR, other exon, 1st Intron, other intron, downstream (<=3kb) and distal Intergenic. We noticed that fruit fly EVEs are most enriched in distal intergenic and intron region, implying that they might function as cis regulatory elements (Figure 3b). To infer the potential functions of those EVEs, we predicted their targets using the “annotatePeak” function from ChIPseeker that regard the nearest gene as the target of corresponding EVE, resulting in 313 putative genes. GO term analysis showed that those putative targets are enriched in GO terms associated with “regulation of organelle organization”, “intracellular signal transduction” and “regulation of vesicle fusion”, “SUMOylation of DNA damage response and repair proteins” and “Transmission across Chemical Synapses” (Figure 3c). As shown in Figure 3d, multiple EVEs elements were located around *step* and *cta* genes, associated with the GO term “regulation of organelle organization”. Further experiments need to be performed to investigate if those EVEs elements indeed regulate the expression patterns of nearby genes.

**Figure 3.**
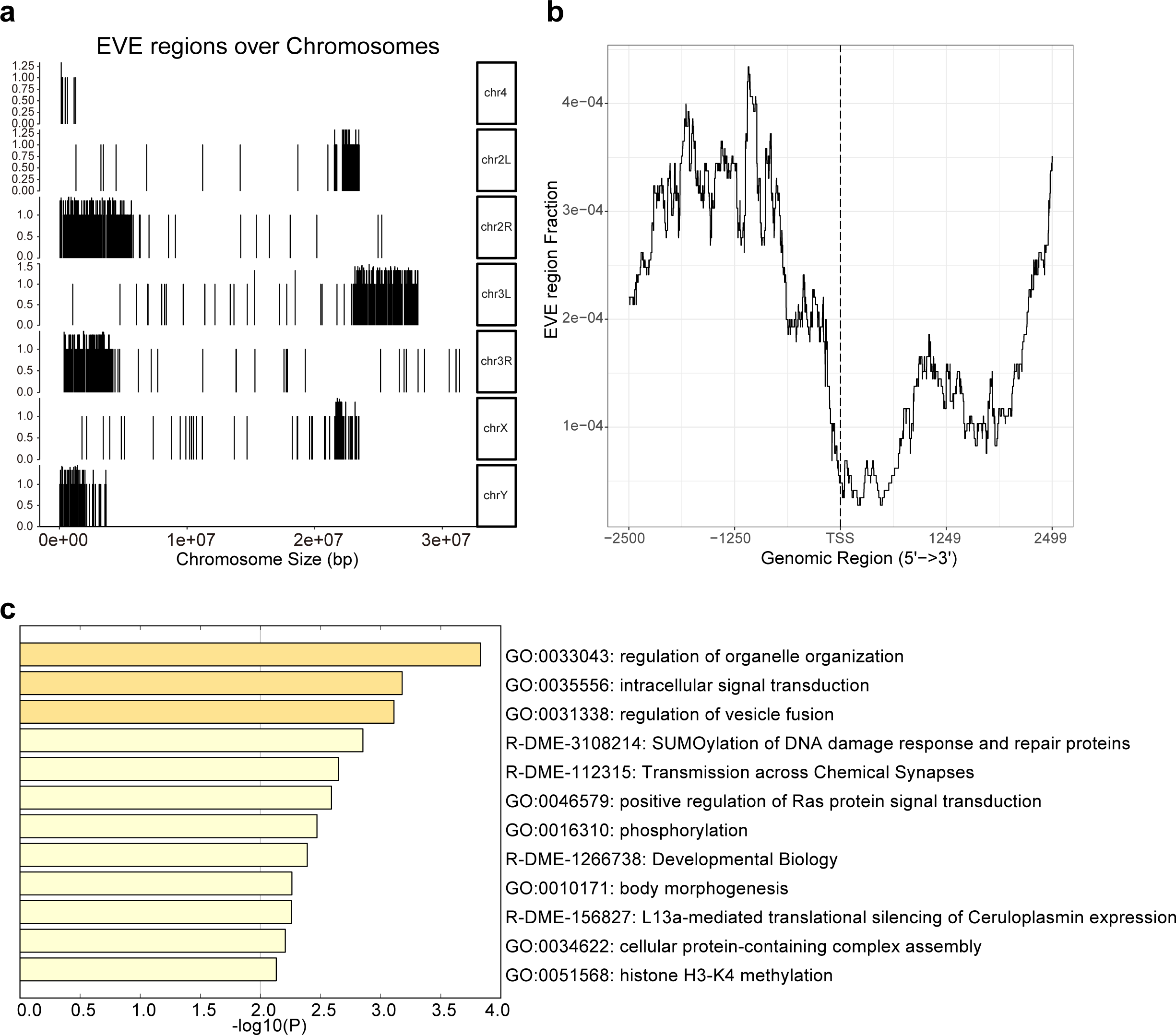

## DISCUSSION

The classification of EVEs from 82 non-mammalian species makes it possible to dissect EVEs in a comprehensive and systematic way. The annotation of EVEs deriving from a variety of literature with a standard script makes the EVEs are comparable to each other which greatly facilitates the comparative analysis on them. Flexible searching options supplied users with powerful and efficient strategies to find EVEs they are interested in. The present of data in user-friendly web interfaces and offering the data for downloading accelerate the sharing and spreading of those valuable EVEs resources. Benefiting from those enormous conveniences provided by FabriEVEs, we observed several unique properties of EVEs in non-mammalian organisms, which give rise to following questions: 1) The copy number of non-retroviral EVEs are much higher in *Phalacrocorax carbo* (cormorant) compared to other bird species. Did it possibly contribute to the evolution of cormorant? 2) The non-retroviral EVEs in birds (225 out of 251) mainly origin from *hepadnaviridae*, while the non-retroviral EVEs reported in invertebrates (136 out of 330) are mainly from *densovirinae*. Is this related to the life styles of *hepadnaviridae* and *densovirinae*? 3) Hundreds of non-retroviruses deriving EVEs were found in fishes, amphibians, birds, reptiles and invertebrates. Of particular interest, high diversity of EVEs were reported in birds and invertebrates. Unlike retroviruses, most non-retroviruses viruses could not integrate into host genome and some RNA viruses do not even have a double strand DNA phase. How did those viruses successfully integrate into animal genome and subsequently become domesticated and co-adapted? 4) In total, 66 copies of complete or nearly full length EVEs were recorded in FabriEVEs. Are those EVEs functional, either working as regulatory elements or actively expressed? To what extends do they differ from their extant counterparts? How could they remain intact after millions of years since the initial integration event. 5) As the progress of genomic, transcriptomics and proteomics studies, there will be plenty of transcription and translation data available for those EVEs. It would be interesting to screen the expression of those EVEs to explore their functions. Overall, those questions illustrate that fascinating biological questions could be inspired by comparing EVEs data sets from various sources integrated into a systematic database such as FabriEVEs.

Though thousands of EVEs in fishes, amphibians, birds, reptiles and invertebrates have been reported, a specialized and dedicated database for them is still missing. FabriEVEs is the first database specially designed to collect, classify, annotate and present non-mammalian EVEs, which will lay the foundation for systematic analysis of EVEs. More importantly, it provides an opportunity to compare EVEs from species with various evolutionary distances, which might speed up our understanding regarding the evolutionary patterns and phylogenic relationships among those species. Currently, there are 1.5 million records in FabriEVEs. We are planning to update our database regularly to include the latest EVEs data reported by researchers in this field. Hopefully, FabriEVEs will help us to get a deeper understanding of the roles EVEs play during the evolution history of non-mammalian animals and throw light upon the co-adaptation between the endogenous viruses and their hosts.

